# Undulating changes in human plasma proteome across lifespan are linked to disease

**DOI:** 10.1101/751115

**Authors:** Benoit Lehallier, David Gate, Nicholas Schaum, Tibor Nanasi, Song Eun Lee, Hanadie Yousef, Patricia Moran Losada, Daniela Berdnik, Andreas Keller, Joe Verghese, Sanish Sathyan, Claudio Franceschi, Sofiya Milman, Nir Barzilai, Tony Wyss-Coray

**Author notes:** Correspondence (T.W-C.) and (B.L.).

## Abstract

Aging is the predominant risk factor for numerous chronic diseases that limit healthspan. Mechanisms of aging are thus increasingly recognized as therapeutic targets. Blood from young mice reverses aspects of aging and disease across multiple tissues, pointing to the intriguing possibility that age-related molecular changes in blood can provide novel insight into disease biology. We measured 2,925 plasma proteins from 4,331 young adults to nonagenarians and developed a novel bioinformatics approach which uncovered profound non-linear alterations in the human plasma proteome with age. Waves of changes in the proteome in the fourth, seventh, and eighth decades of life reflected distinct biological pathways, and revealed differential associations with the genome and proteome of age-related diseases and phenotypic traits. This new approach to the study of aging led to the identification of unexpected signatures and pathways of aging and disease and offers potential pathways for aging interventions.

---

Aging underlies progressive changes in organ functions and is the primary risk factor for a large number of human diseases^1^. A deeper understanding of aging is thus likely to provide insight into the underlying mechanisms of disease and facilitate the development of novel therapeutics that target the aging process more generally. A growing number of investigators have applied genomic, transcriptomic and proteomic assays (collectively referred to as omics) to studies of aging from model organisms to humans^2^. Human genetic studies have uncovered relatively few modifiers of aging, yet other omics modalities, which measure more dynamic gene modifications or products, have provided valuable insight. For example, the transcriptome varies greatly during aging across tissues and organisms^3^, pointing to evolutionarily conserved, fundamental roles of developmental and inflammatory pathways^4^. Protein composition of cells, body fluids, and tissues change similarly with age and provide insight into complex biological processes since proteins are often direct regulators of cellular pathways. In particular, blood, which contains proteins from nearly every cell and tissue, has been extensively analyzed to discover biomarkers and gain insight into disease biology. Accordingly, organismal aging results in proteomic changes in blood that reflect aspects of aging of different cell types and tissues.

Perhaps the strongest evidence that blood can be used to study aging comes from experiments employing heterochronic parabiosis, a surgically induced state in which the circulatory systems of young and old mice are joined. These studies show that multiple tissues, including muscle, liver, heart, pancreas, kidney, bone, and brain can be rejuvenated in old mice^5–13^. Plasma (the soluble fraction of blood) from old mice is sufficient to accelerate aspects of brain aging following repeated infusion into young mice^12^ and young plasma can slow and reverse memory impairment and other aspects of brain aging^13,14^. Altogether, these studies support the notion that the plasma proteome harbors key regulators of aging. Identifying such protein signatures may help understand mechanisms of organismal aging. However, blood proteomic changes with age have not been thoroughly exploited and require novel tools to derive insight into the biology of aging. Here, we carried out a deep proteomic analysis of plasma from young adults to nonagenarians and applied newly developed data analysis tools. We discovered distinct changes in protein expression across stages of lifespan and link these changes to biological pathways and disease.

## Results

### Linear modeling links the plasma proteome to functional aging and identifies a conserved aging signature

We analyzed plasma isolated from ethylenediaminetetraacetic acid (EDTA)-treated blood acquired via venipuncture from 4,331 healthy individuals aged 18-95 years from the Interval and LonGenity cohorts (Fig. 1a, Supplementary Fig. 1). Currently, one of the most advanced tools for the measurement of plasma proteins are single-stranded oligonucleotides known as aptamers^15,16^, which bind their target with high affinity and specificity. To generate a proteomic dataset of human lifespan, we used the SomaScan aptamer technology, capable of quantifying thousands of proteins involved in intercellular signaling, extracellular proteolysis, and metabolism (Supplementary Tables 1 and 2). Reproducibility analysis based on technical replicates showed an overall low median coefficient of variation (CV) of ~5% within runs and 10% between runs^17^ (Supplementary Table 3). The Interval and LonGenity proteomics datasets analyzed in this study can be interrogated by an interactive web interface (https://twc-stanford.shinyapps.io/aging_plasma_proteome/).

**Figure 1:**
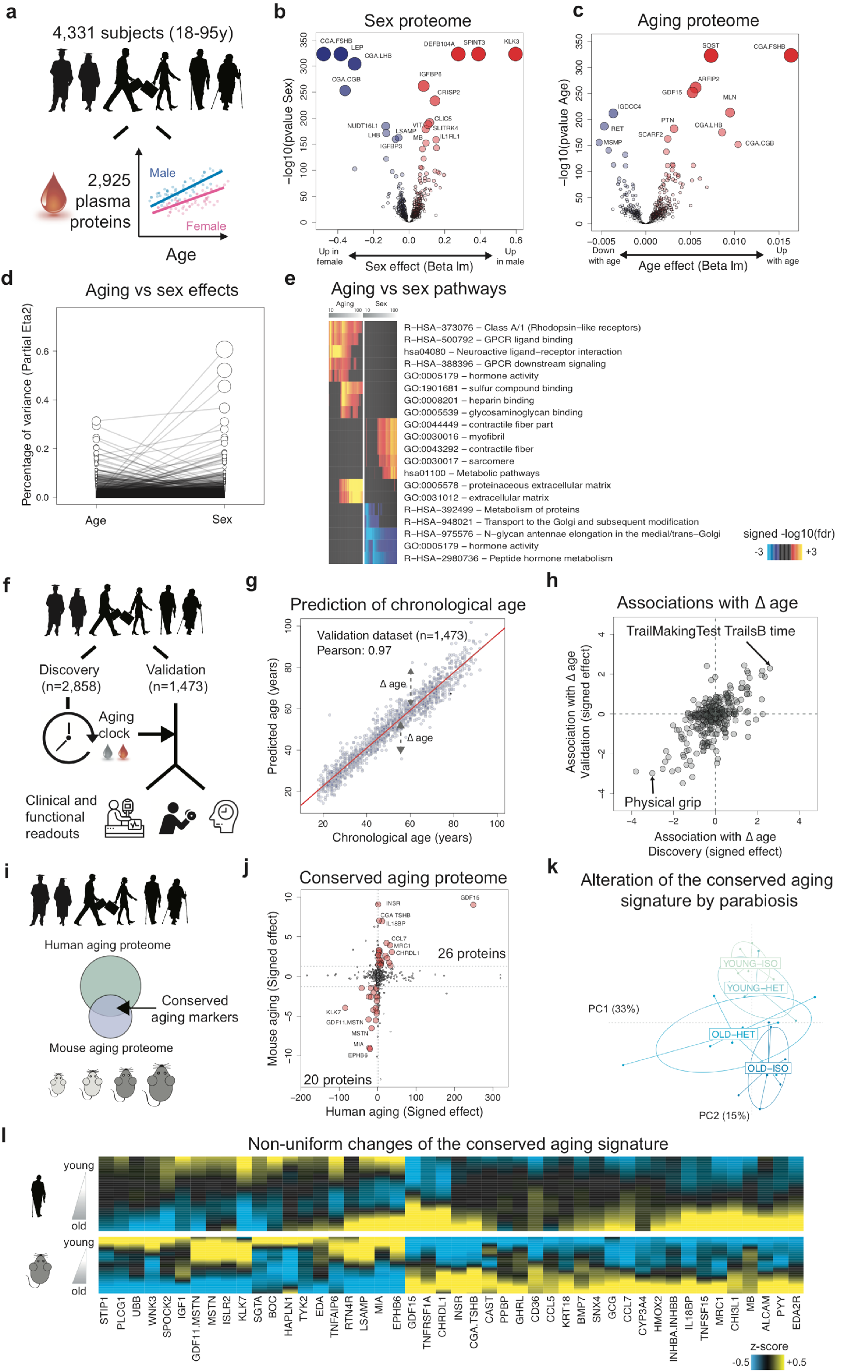
Linear modeling links the plasma proteome to functional aging and identifies a conserved aging signature. (a) Schematic representation of the linear analysis of the plasma proteome. Four thousand three hundred thirty-one human EDTA plasma samples were measured using the SomaScan platform and analyzed using linear models, adjusted for age, sex and subcohort. Volcano plots representing changes of the plasma proteome with sex (b) and age (c). Dot sizes are proportional to the product of −log10(p-value) and sex or age effect (beta of the linear model). (d) Relative percentage of variance explained by age and sex. Partial Eta2 is calculated for age and sex. Values for each plasma protein are connected by edges. (e) Pathways associated with sex and age identified by Sliding Enrichment Pathway Analysis (SEPA). Enrichment for pathways in the top 10 to top 100 proteins ranked by the product of - log10(p-value) and beta lm is tested, in increments of 1 protein. Proteins upregulated and downregulated are analyzed separately. The top 10 pathways per condition are represented. (f) Modeling chronological age using the plasma proteome analysis. A LASSO model with 10-fold cross validation was trained on 2/3 of the subjects (n=2,858 out of 4,331). The model was tested on the remaining 1,473 samples. Then, association between delta age (difference between predicted age and chronological age) with functional readouts was investigated. (g) Prediction of age in the validation cohort (n=1,473) using 373 plasma markers. Pearson correlation coefficient between observed and predicted age is indicated. (h) Association between delta age (difference between observed age and chronological age) and functional readouts in old. Linear modeling between 334 functional readouts and delta age adjusted for age at clinical visit, sex and cohort was tested. Top associations in both discovery and validation datasets are represented. (i) Schematic representation of the comparison between the human and mouse aging proteomes. (j) Conserved markers of aging. Effect of aging on the human plasma proteome is compared to the effect of aging on the mouse plasma proteome. Both human and mouse aging effects are signed by the beta age of their corresponding linear analysis. Forty-six plasma proteins are changing in the same direction in mouse and humans (red dots) and define a conserved aging signature. (k) Alteration of the conserved aging signature by parabiosis. Normed principal component analysis was used to characterize changes of the conserved aging signature when mice are exposed to young or old blood. (l) Age-related changes of the conserved aging signature. Plasma protein levels were z-scored and aging trajectories were estimated by locally estimated scatterplot smoothing (LOESS).

Since females have longer average lifespans than males independent of socio-economic status^18^, we sought to determine whether sex and aging proteomes are interconnected (Fig 1b-1d). Proteins most strongly changed with sex included well-known follicle stimulating hormone (CGA FSHB), human chorionic gonadotropin (CGA CGB), and prostate-specific antigen (KLK3). With age, the most prominent overall changes with respect to fold change and statistical significance included sclerostin (SOST), ADP Ribosylation Factor Interacting Protein 2 (ARFIP2), and growth differentiation factor 15 (GDF15), in addition to several sex proteins such as CGA FSHB. The proteins most strongly associated with age also changed significantly with sex, (Fig. 1d); 895 proteins out of the 1,379 proteins altered with age (q<0.05) were significantly different between males and females (q<0.05, Supplementary Table 4). These results are aligned with a growing number of studies demonstrating that males and females age differently^19^. To determine whether these findings are representative of the general population, we compared changes identified in this study with findings from 4 small independent cohorts (n=171, age range 21-107y, Supplementary Fig. 1d) and with findings from an independent study^20^. Even though these independent studies used an older version of the SomaScan assay measuring only a subset of the current proteins (1,305 proteins; Supplementary Table 2), we observed high consistency of the aging and sex proteomes across cohorts (Supplementary Fig. 2).

To establish the biological relevance of these changes, we queried GO, KEGG and Reactome databases and measured enrichment of proteins in pathways using a “Sliding Enrichment Pathway Analysis (SEPA)” (Supplementary Tables 5 and 6, see Methods). The heatmaps produced by SEPA firstly illustrate the relationship between the top 100 proteins and the biological pathways they represent; secondly, the heatmaps emphasize how a restricted list of top aging proteins reveal biological pathways which would have escaped common pathway mining modalities. SEPA indicated that incremental lists of proteins are needed to determine biological functions of sex proteins, and pointed to expected differences in hormonal metabolism and activity (Fig. 1e, Supplementary Table 6). Conversely, an extensive list of aging proteins identified enrichment for blood-related pathways such as heparin and glycosaminoglycan binding, as recently reported^20^.

To determine whether the plasma proteome can predict chronological age and serve as a “proteomic clock,” we used 2,858 randomly selected subjects to fine-tune a predictive model that was tested on the remaining 1,473 subjects (Fig. 1f). We identified a sex-independent plasma proteomic clock consisting of 373 proteins (Supplementary Table 7). This clock was highly accurate in predicting chronological age in the discovery, validation and 4 small independent cohorts (r=0.93-0.97, Fig. 1g and Supplementary Fig. 3a-b). Remarkably, subjects that were predicted younger than their chronologic age based on their plasma proteome performed better on cognitive and physical tests (Fig. 1h and Supplementary Table 8). While a reduced model comprising only 9 proteins predicted age with good accuracy (Supplementary Fig.3c and Supplementary Table 7), a combination of different sets of proteins may be required to model changes in a large set of clinical and functional parameters (Supplementary Fig.3d).

Since most biological pathways that change with age are evolutionarily conserved from yeast to mammals^21^, we tried to identify possible aging proteins conserved between mice and humans. Thus, we analyzed mouse plasma (n=110, aged 1 month to 30 months) using the SomaScan assay, which reliably measure hundreds of proteins in samples from non-human species^22^ and has proven useful in some applications in recent mouse studies^9,23^ (Fig. 1i, Supplementary Table 9). In mice, 172 proteins changed with age (out of 1,305 measured, Supplementary Table 10, q<0.05) and 46 proteins overlapped with human aging proteins (Fig. 1j). Remarkably, many of these proteins were modulated by heterochronic parabiosis: young mice exposed to old plasma (young-heterochronic mice) showed a relatively older plasma signature while aged mice exposed to young plasma (old-heterochronic mice) showed a younger signature (Fig. 1k). Altogether, standard linear modeling of the plasma proteome during human lifespan revealed established aging pathways, possibly indicating accelerated and decelerated aging in humans and mice. Intriguingly, changes to the conserved aging proteins did not occur simultaneously (Fig. 1l). Thus, the chronology of aging in the plasma proteome requires further investigation.

### Clustering individual protein trajectories reveals undulation of the aging plasma proteome

While the above standard linear modeling showed prominent overall changes in plasma protein composition, we were struck by the undulating behavior of the 46 conserved proteins (Fig. 1l) and more globally by the 2,925 plasma proteins as a group when they were visualized as z-scored changes across lifespan (Fig. 2a-b). These undulating patterns were detected in independent human cohorts and in mice (Supplementary Fig. 4), suggesting they are robust and conserved across species.

**Figure 2:**
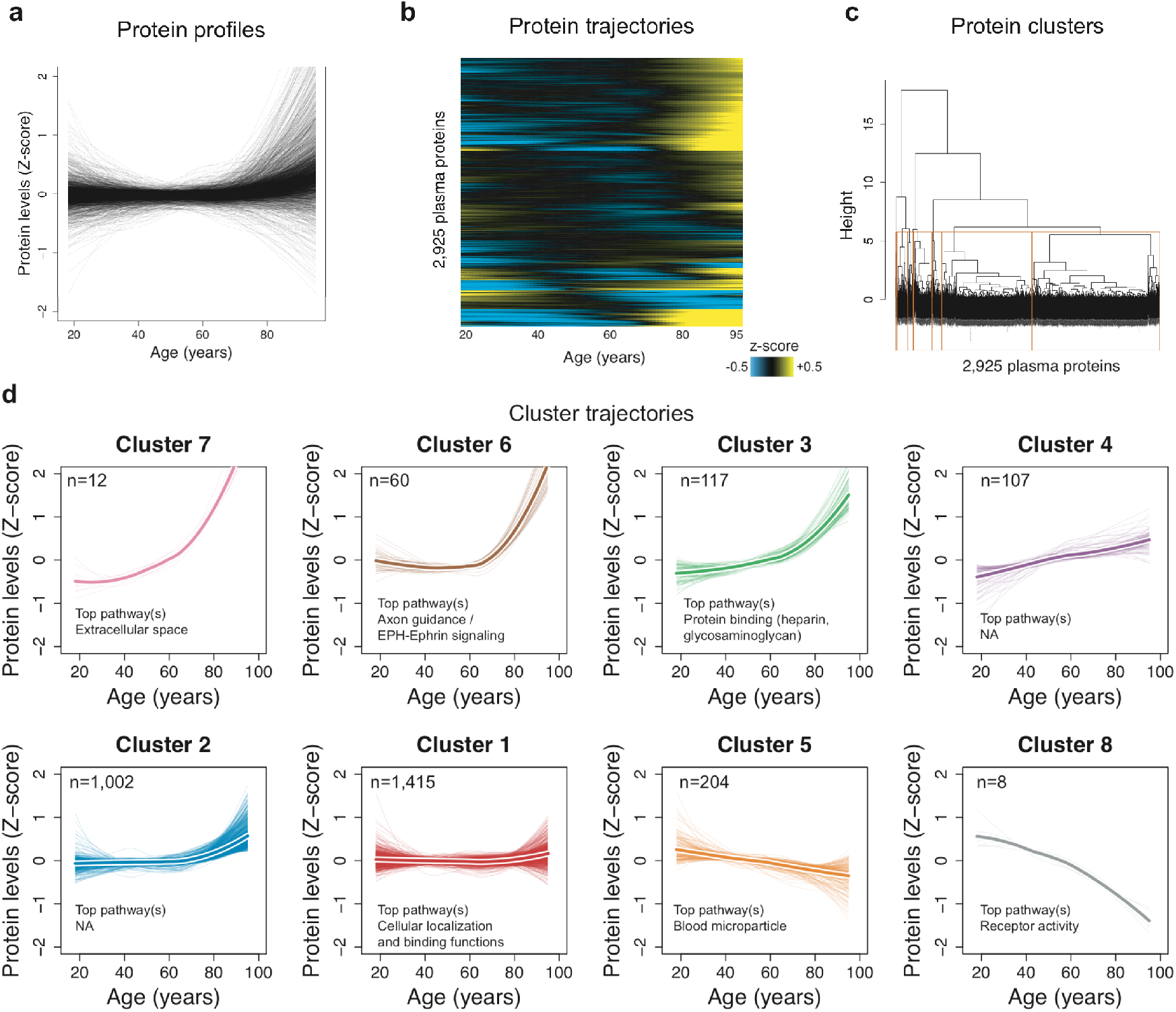
Clustering of protein trajectories identifies linear and non-linear changes during aging. (a) Protein trajectories during aging. Plasma protein levels were z-scored and trajectories of the 2,925 plasma proteins were estimated by LOESS. (b) Trajectories are represented in two dimensions by a heatmap and unsupervised hierarchical clustering was used to group plasma proteins with similar trajectories. (c) Hierarchical clustering dendrogram. The 8 clusters identified are represented by orange boxes. (d) Protein trajectories of the 8 identified clusters. Clusters are grouped by the similarity of global trajectories, the thicker lines representing the average trajectory for each cluster. The number of proteins and the most significant enriched pathways are represented for each cluster. Pathway enrichment was tested using GO, Reactome and KEGG databases. The top pathways are indicated and the top 20 pathways for each cluster are listed in supplementary table 12.

To reduce complexity of these changes, we grouped proteins with similar trajectories using unsupervised hierarchical clustering (Fig. 2c) and identified 8 clusters of protein trajectories changing with age, which ranged in size from 8 to 1,415 proteins (Supplementary Table 11). In addition to linear trajectories (clusters 1, 4 and 5), several non-linear trajectories including stepwise, logarithmic and exponential changes were also evident (clusters 2, 3, 6, 7 and 8) (Fig. 2d). Remarkably, these cluster trajectories were similarly detectable in independent cohorts (Supplementary Fig. 5). Out of the 8 clusters analyzed, 6 were enriched for specific biological pathways (q<0.05; Supplementary Fig. 6, Supplementary Table 12), suggesting distinct, yet orchestrated changes in biological processes during the human lifespan. For example, proteins present in blood microparticles consistently decreased with age (cluster 5); other blood-related pathways such as heparin and glycosaminoglycan binding increased in a two-step manner (cluster 4); while levels of proteins involved in axon guidance and EPH-ephrin signaling remain constant until age 60 before they rise exponentially (cluster 6) (Fig. 2d). Altogether, we find that the majority of changes in the plasma proteome during lifespan occur in a non-linear manner.

### Quantification of proteomic changes across human lifespan uncovers waves of aging proteins

To gain a quantitative understanding of the proteomic changes occurring throughout human lifespan, we developed the software tool Differential Expression - Sliding Window ANalysis (DE-SWAN) (Fig. 3a). This algorithm analyzes levels of plasma proteins within a window of 20 years and compares two groups of individuals in parcels of 10 years (e.g. 35–45 compared with 45–55). We used a window of 20 years to detect age related-changes and slid the window in increments of 1 year from young to old. By this method, we aimed to detect changes at particular stages of lifespan and determine the sequential effects of aging on the plasma proteome (while also controlling for the effect of confounding factors). This approach identified hundreds of plasma proteins changing in waves throughout age (Fig. 3b). Summing the number of differentially expressed proteins at each age uncovered 3 crests at ages 34, 60, and 78 (Fig. 3c, Supplementary Fig. 7a and Supplementary Table 13). These crests disappeared when the ages of individuals were permutated (Supplementary Fig. 7b), but were still detectable when using different statistical models (e.g. smaller/larger sliding windows) (Supplementary Fig. 7c, and 8, Supplementary Table 13), indicating robustness of these age-related waves of plasma proteins.

**Figure 3:**
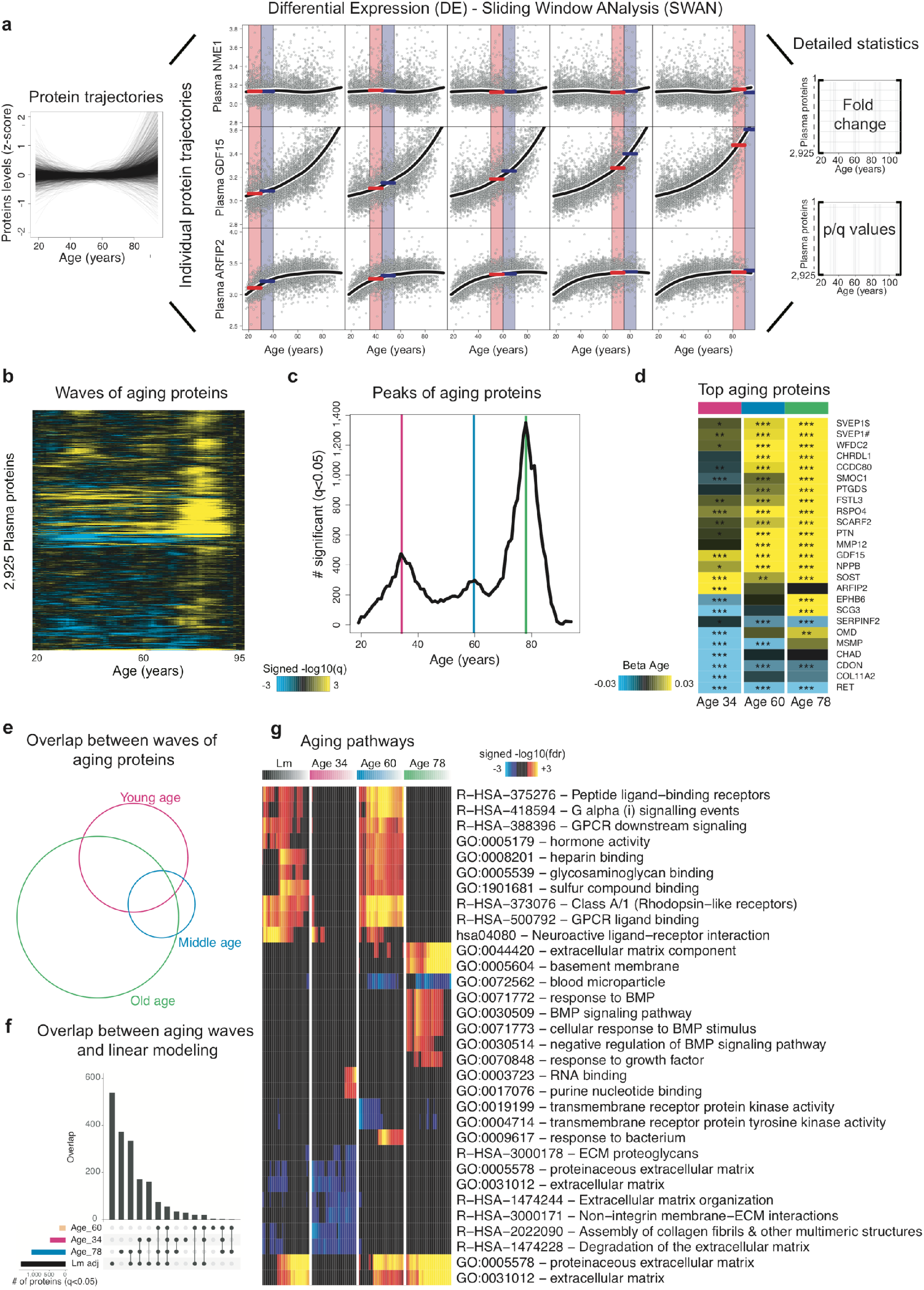
Sliding window analysis distinguishes waves of aging plasma factors. (a) Principles of Differential Expression Sliding Window Analysis (DE-SWAN) compares levels of plasma proteins between groups of individuals in parcels of 10 years, e.g. 30–40 compared with 40–50. By sliding the window, DE-SWAN scans the whole aging plasma proteome and identifies linear and non-linear changes with age. Examples of the DE-SWAN approach for 3 plasma proteins and 5 age windows are represented. Red and blue rectangles show the two parcels that are compared and the red and blue lines symbolize the mean within each parcel. DE-SWAN provides statistics for each age window and each plasma protein, allowing a detailed analysis of plasma proteomic changes during aging. (b) Waves of aging plasma proteome characterized by DE-SWAN. Within each window, −log10(p-values) and −log10(q-values) were estimated by linear modeling adjusted for age and sex. Local changes attributable to age were signed based on corresponding beta age. (c) Number of plasma proteins differentially expressed during aging. Three local peaks at the ages of 34, 60 and 78 were identified by DE-SWAN. (d) Top 10 plasma proteins identified by linear modeling and DE-SWAN at age 34, 60 and 78. Blue and yellow colors represent local increase and decrease, respectively. # and $ indicate different SOMAmers targeting the same protein. * q<0.05, ** q<0.01, *** q<0.001 (e) Intersections between aging plasma proteins identified by linear modeling and DE-SWAN at ages 34, 60 and 78 (q<0.05) visualized by Venn diagram. (f) Intersections between the waves and linear modeling (q<0.05) visualized by Upset plot. (g) Visualization of pathways significantly enriched for aging proteins identified by linear modeling and DE-SWAN at age 34, 60 and 78. Enrichment in the top 10 to top 100 proteins were tested in increments of 1 protein. Proteins upregulated and downregulated were analyzed separately. GO, Reactome and KEGG databases are shown. The top 10 pathways per condition are represented.

The 3 age-related crests were comprised of different proteins (Fig. 3d, Supplementary Table 14). Few proteins, such as GDF15, were among the top 10 differentially expressed proteins in each crest, consistent with its strong increase across lifespan (Fig. 3a). Other proteins, like chordin-like protein 1 (CHRDL1) or pleiotrophin (PTN), were significantly changed only at the last two crests, reflecting their exponential increase with age. The overlap between the sets of proteins changing at age 34, 60 and 78 was statistically significant (p<0.05) but limited (Fig. 3e) and most of the proteins changing in old age were not identified by linear modeling (Fig. 3f). This prompted us to use SEPA to determine whether these waves reflected distinct biological processes. Strikingly, we observed a prominent shift in multiple biological pathways with aging (Fig. 3g). At young age (34 years), we observed a downregulation of proteins involved in structural pathways such as the extracellular matrix (ECM). These changes were reversed in middle and old ages (60 and 78 years, respectively). At age 60, we found a predominant role of hormonal activity, binding functions and blood pathways. At age 78, key processes still included blood pathways but also bone morphogenetic protein signaling, which is involved in numerous cellular functions, including inflammation^24^. Pathways changing with age according to linear modeling (Fig. 1g) overlapped most strongly with the crests at age 34 and 60 (Fig. 3f), indicating the dramatic changes in protein expression occurring in the elderly might be masked in linear modelling by more subtle changes at earlier ages. Altogether, these results suggest that aging is a dynamic, non-linear process characterized by waves of changes in plasma proteins that are reflective of a complex shift in the activity of biological processes.

### Proteins linked to age-related diseases are enriched in distinct waves of aging

The plasma proteome is sensitive to the physiological state of an individual but is also genetically determined^25^. To deconvolute this complexity between genome, proteome, and physiology, we asked whether the top plasma aging proteins change their levels due to genetic polymorphisms (pQTLs) or whether they are among the top predictors of disease, or phenotypic traits. More specifically, we sought to determine whether proteins that comprised the 3 waves of aging were uniquely linked to the genome or proteome of age-related diseases and traits (Fig. 4a). To this end, we used the ranked lists of the top proteins identified by DE-SWAN at each of the three crests (Fig. 3c and Supplementary Table 14) and summed the number of proteins linked to the genome and proteome of specific diseases and traits separately for each wave (i.e. the cumulative sum) (Fig. 4b-i and Supplementary Fig. 9). First, we mined the genomic atlas of the human plasma proteome^25^ (Fig. 4b) and discovered that the aging proteome is also genetically determined (Fig.4c and Supplementary Fig. 9). However, the rank of the proteins determined by trans-association appeared more random with aging (Fig. 4c), suggesting that other sources are driving the aging plasma proteome with age. We then tested whether the waves of aging proteins were differentially linked with changes in cognitive and physical functions identified in Fig. 1h. Interestingly, the proteome associated with these traits overlapped with the proteome defining middle and old ages, i.e. when these functions decline the most (Fig. 4d and Fig. 4e). Finally, we used public datasets and summary statistics from SomaScan proteomic studies focusing on age-related diseases including Alzheimer’s Disease (AD)^26^, Down Syndrome (DS)^27^ and cardiovascular disease (CVD)^28^. A proteomic study predicting body mass index (BMI)^29^ based on the plasma proteome was used as a control since weight gain varies widely with age according to data from the US National Health and Nutrition Examination Study^30^ (Supplementary Fig. 10). As expected, the proteome linked to BMI was not selectively enriched for proteins defining waves of aging (Fig. 4f). CVD-associated proteins were strongly enriched in waves of proteins defining middle and old age compared to young age (Fig. 4g). This enrichment corresponded to an increased incidence of CVD after 55 years^31^. Finally, AD- and DS-associated proteins overlapped with the top proteins defining middle age and old age but not with proteins in the young wave of aging (Fig. 4h-i). The fact that the proteome defining these two diseases also changed in old individuals of a separate disease-free cohort supports the notion of accelerated aging in DS and AD^32,33^. Altogether, these results show that the waves of proteomic aging are differentially linked to the genomic and proteomic traits of various diseases.

**Figure 4:**
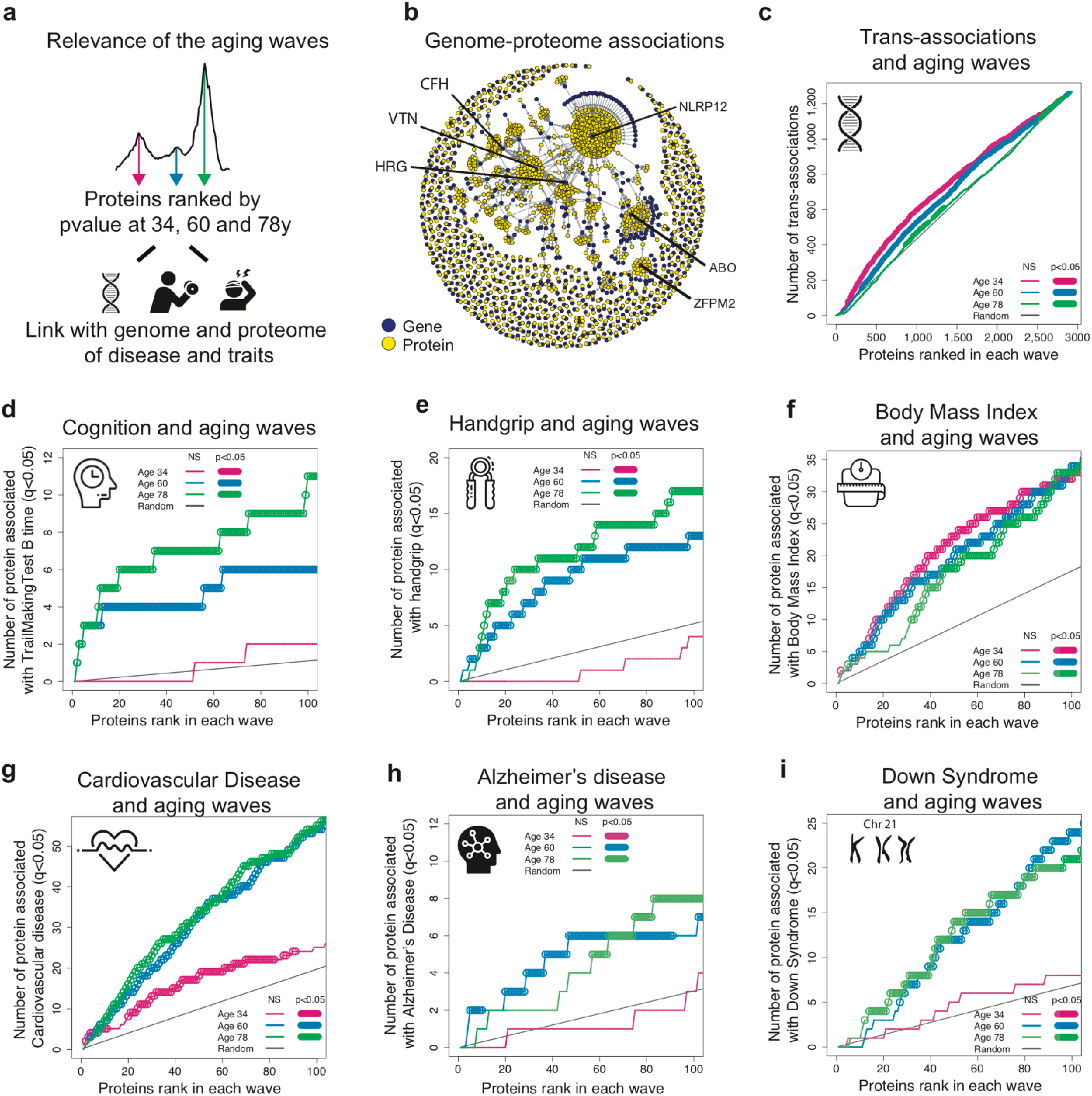
Waves of aging proteins are differentially linked to the genome and proteome of disease and traits. (a) Relevance of the aging waves. Schematic representation of analysis. The proteins changing at 34, 60 and 78y were ranked by p-value and were associated with the genome and the proteome of disease and traits. (b) Association between the genome and the proteome. Network created using the pQTL associations identified by Sun et al. (2018). (c) Enrichment for trans-association in the waves of aging proteins identified by DE-SWAN. Aging proteins at age 34, 60 and 78y were ranked based on p-value and the cumulative number of transassociations was enumerated. Permutation based tests (1e+5) were used to assess significance. Enrichment for proteins involved in cognitive and physical performance in the waves of aging proteins identified (d-e). Enrichment for disease-associated proteins in the waves of aging proteins (f-i).

## Discussion

Our analysis of 2,925 proteins in plasma from healthy humans reveals complex, non-linear changes during human lifespan. Although modeling protein trajectories is necessary to truly appreciate these undulating changes, a standard linear analysis provides key information about the aging plasma proteome.

It is well known that men and women age differently^19^, but we were surprised to find that 2/3 of the proteins changing with age are also changing with sex (895 proteins out of the 1,379 proteins changing with age). These results strongly support the National Institutes of Health (NIH) policy on the inclusion of women in clinical research and the inclusion of sex as a biological variable in scientific experiments. Nevertheless, a unique proteomic clock can be used to predict age in men and women and deviations from this plasma proteomic clock are correlated with changes in clinical and functional parameters (Fig 1g-h). The panel of 373 proteins or a subset can be used to assess the relative health of an individual and to measure healthspan, analogous to epigenetic clocks based on DNA methylation patterns^34^. More large-scale plasma proteomic studies will be required to establish the validity and utility of this clock and to test if specific subsets of proteins may be more appropriate to reflect particular clinical and functional parameters.

Blood is a sensitive marker of functional aging but also plays an active role in aging. Numerous studies have shown that soluble factors from young mouse blood reverse aspects of aging and disease across multiple tissues^5–13,35^. Here, we describe a 46-protein aging signature that is conserved in humans and mice, containing several known aging proteins such GDF15^36^ and IGF1/INSR^37^ but also less investigated ones (Fig 1l). This conserved signature will allow deeper investigation of translational aging interventions in mice, such as heterochronic parabiosis, which partially reverses age-related changes of these proteins (Fig 1k).

By deep mining the aging plasma proteome, we identify undulating changes during the human lifespan. These changes are the result of clusters of proteins moving in distinct patterns, culminating in the emergence of 3 waves of aging. Somewhat unexpectedly, we find that these clusters are often part of shared biological pathways, most notably in cellular signaling functions (Fig. 2). In addition, we identify and provide biological relevance for the 3 main waves of aging proteins (Fig. 3). These waves are characterized by key biological pathways with little overlap, demonstrating changes to chronological aging in the plasma proteome. In comparison, linear modeling fails to identify changes occurring late in the 8^th^ decade of life (Fig. 3g). We conclude that linear modeling of aging based on “omics” data does not capture the complexity of biological aging across organismal lifespan. Thus, DE-SWAN will be invaluable for analyzing longitudinal or cross-sectional datasets with non-linear, quantifiable changes, and for integrating non-linear changes in the analysis of high dimensional “omics” datasets of a quantitative trait.

Sources of variation of the plasma proteome can be diverse and a majority of the plasma proteome is under substantial genetic control^25,38^. Intriguing, we observed that the relative importance of trans-associations decreased with aging (Fig. 4c). This led us to investigate other sources of variance with a focus on disease-associated proteomes and traits. We found that proteins comprising the middle and old age waves differentially overlapped with proteins associated with cognitive and physical impairments. Moreover, plasma proteins that changed most prominently with age also discriminated patients from age-matched controls in age-related diseases including AD, DS and CVD (Fig. 4g-i). This suggests that the characteristic plasma proteins of aging are amplified in age-related diseases. Using an AD- and DS-free cohort, we provide evidence to support the concept of accelerated aging for these two diseases^32,33^. Further investigation of these proteins is warranted to determine whether these associations indicate aging biomarkers and/or are causal proteins of disease. Nonetheless, these results suggest that variance within the aging plasma proteome slowly transitions from hard coding factors (i.e. genomic) to soft coding factors (e.g. diseases, environmental factors and resulting changes in cognitive and physiological functions).

The undulating nature of the aging plasma proteome and its interactions with diseases ought to be considered when developing proteomic signatures for diagnostic purposes. Indeed, disease proteomes overlap significantly with the waves of aging proteins (Supplementary Table 15). Taking into account the heterogeneous and non-linear changes of the plasma proteome during the entire lifespan can likely improve the sensitivity and specificity of prognostics and diagnostics tests. Moreover, these results are pertinent when considering the use of blood or blood products to treat aging and age-related diseases^39^. Specifically, identifying proteins within plasma that promote or antagonize aging at different stages of life could lead to more targeted therapeutics, as well as preventative therapies. Such reliable tests and treatments are still urgently needed for numerous diseases and, in the future, we hope to describe plasma proteome changes that predict subjects transitioning to disease. Of particular interest are studies of AD for which blood-based biomarkers are unavailable and clinical symptoms are believed to occur up to two decades after disease onset.

### URLs

https://twc-stanford.shinyapps.io/aging_plasma_proteome/

### Online content

Any methods, additional references, Nature Research reporting summaries, source data, statements of data availability and associated accession codes are available at https://doi.org/xxxxyyyy.

## Acknowledgements

We thank the members of the Wyss-Coray laboratory for feedback and support. We thank clinical staff for human blood-plasma collection/coordination. We thank Adam Butterworth for his help to get access to the Interval proteomics data.

The AddNeuroMed data are from a public-private partnership supported by EFPIA companies and European Union of the Sixth Framework program priority FP6-2004-LIFESCIHEALTH-5. Clinical leads responsible for data collection are Iwona Kłoszewska (Lodz), Simon Lovestone (London), Patrizia Mecocci (Perugia), Hilkka Soininen (Kuopio), Magda Tsolaki (Thessaloniki), and Bruno Vellas (Toulouse) and imaging leads Andy Simmons (London), Larsw Olad Wahlund (Stockholm) and Christian Spenger (Zurich) and bioinformatics leads are Richard Dobson (London) and Stephen Newhouse (London).

This work was supported by a National Institutes of Health National Institute on Aging (NIA) F32 1F32AG055255 01A1 (D.G.), the Cure Alzheimer’s Fund (T.W-C.), the NOMIS Foundation (T.W-C.), the Stanford Brain Rejuvenation Project (an initiative of the Stanford Wu Tsai Neurosciences Institute), The Paul F. Glenn Center for Aging Research (T.W-C.), NIA R01 AG045034; DP1 AG053015 (T.W-C.) and the NIA funded Stanford Alzheimer’s Disease Research Center P50AG047366, NIA K23AG051148 (S.M.), R01AG061155 (S.M.), American Federation for Aging Research (S.M.), R01AG044829 (J.V. and N.B), NIA R01AG057909 (N.B.), the Nathan Shock Center of Excellence for the Basic Biology of Aging P30AG038072 (N. B.), the Glenn Center for the Biology of Human Aging (N.B.).

## Author Contributions

B.L. and T.W-C. planned the study, D.B., C.F., S.M., J.V., S.S. and N.B. provided human plasma samples, N. S., S.E.L and H.Y performed the mouse experiments, B.L analyzed the data with contributions from T.N. and A.K., P.M.L. developed the shiny app, B.L., D.G. and T.W-C. wrote the manuscript, A.K, C.F, S.M, J.V., S.S., N.B. and T.W-C. supervised the study, all authors edited and reviewed the manuscript.

## Additional Information

**Supplementary information** is available for this paper at https://doi.org/xxxxyyyt.

**Reprints and permissions information** is available at www.nature.com/reprints.

**Correspondence and requests for materials** should be addressed to T.W-C or B.L.

**Publisher’s note:** Springer Nature remains neutral with regard to jurisdictional claims in published maps and institutional affiliations.

## Methods

### Plasma proteomics measurements

The SomaScan platform was used to quantify relative protein levels. This platform was established to identify biomarker signatures of diseases and conditions, including cardiovascular risk^28^, cancer^40^ and neurodegenerative diseases^26^. The SomaScan platform is based on modified single-stranded DNA aptamers (SOMAmer^®^ reagents) binding to specific protein targets. Assay details have been previously described^17^. Different versions of the SomaScan assay were used in the LonGenity, INTERVAL and the 4 independent human cohorts. These versions contained 5,284, 4,034, and 1,305 aptamers, respectively.

Out of the 4,034 aptamers measured in the INTERVAL cohort, 3,283 were contained in the publicly available dataset (https://ega-archive.org/studies/EGAS00001002555). Our study focuses on 2,925 aptamers with identical SeqId and SeqIdVersion in both INTERVAL and LonGenity cohorts (Supplementary Table1). Out of the 2,925 aptamers, 888 were measured in the 4 independent cohorts and in mice (Supplementary Table 2).

### Human cohort characteristics

#### INTERVAL cohort

Participants in the INTERVAL randomized controlled trial were recruited with the active collaboration of the National Health Service Blood and Transplant England (www.nhsbt.nhs.uk), which has supported field work and other elements of the trial. DNA extraction and genotyping was co-funded by the National Institute for Health Research (NIHR), the NIHR BioResource (http://bioresource.nihr.ac.uk/) and the NIHR Cambridge Biomedical Research Centre at the Cambridge University Hospitals NHS Foundation Trust. The INTERVAL study was funded by NHSBT (11-01-GEN). The academic coordinating center for INTERVAL was supported by core funding from: NIHR Blood and Transplant Research Unit in Donor Health and Genomics (NIHR BTRU-2014-10024), UK Medical Research Council (MR/L003120/1), British Heart Foundation (RG/13/13/30194) and the NIHR Cambridge Biomedical Research Centre at the Cambridge University Hospitals NHS Foundation Trust. Proteomic assays were funded by the academic coordinating center for INTERVAL and MRL, Merck & Co., Inc. A complete list of the investigators and contributors to the INTERVAL trial has been previously reported^41^. The academic coordinating center would like to thank blood donor center staff and blood donors for participating in the INTERVAL trial.

Proteomics measurements from 3,301 human plasma samples (1,685 males and 1,616 females) from 2 different subcohorts were used for this study. Age ranged from 18 to 76 years with a median age of 45 (1^st^ Quartile=31, 3^rd^ Quartile=55). Sample selection, processing and preparation were detailed previously^25^.

#### LonGenity cohort

LonGenity is an ongoing longitudinal study initiated in 2008, designed to identify biological factors that contribute to healthy aging^42^. The LonGenity study enrolls older adults of Ashkenazi Jewish descent, age 65-94 years at baseline. Approximately 50% of the cohort consists of offspring of parents with exceptional longevity (OPEL), defined by having at least one parent that survived to 95 years of age. The other half of the cohort includes offspring of parents with usual survival (OPEL), who do not have a parental history of exceptional longevity. Proteomics measurements from 1,030 human plasma samples (457 males and 573 females) collected at baseline in LonGenity participants were used for this study. Age ranged from 61 to 95 years with a median age of 74 (1^st^ Quartile=69, 3^rd^ Quartile=80). LonGenity subjects are thoroughly characterized demographically and phenotypically at annual visits that include collection of medical history and physical and neurocognitive assessments. Sixty-eight subjects without clinical and functional data were excluded from the analysis. The LonGenity study was approved by the Institutional Review Board (IRB) at the Albert Einstein College of Medicine.

#### Additional 4 small independent cohorts

One hundred seventy-one human plasma samples (84 males and 87 females) were obtained from 4 different cohorts (VASeattle, PRIN06, PRIN09, GEHA). Age ranged from 21 to 107 years with a median of 70 years (1^st^ Quartile=58, 3^rd^ Quartile=89). Written informed consent was obtained for each subject. The IRB has determined that our research does not meet the definition of human subject research per Stanford’s Human Research Protection Program policy and no IRB approval was required for this study.

For these cohorts, all samples were stored at −80 °C and 150μl aliquots of plasma were sent on dry ice to SomaLogic Inc. (Boulder, Colorado, US). Plasma samples were analyzed in three different batches, 24 samples in 2015, 70 in 2016 and 77 in 2017. In addition to these 171 plasma samples, 12 additional aliquots from 4 of these samples were measured in the different batches to estimate intra- and inter-assay variability (Supplementary Table 3). Data for 1,305 SOMAmer probes were obtained. No sample or probe data were excluded. HybNorm.plateScale.medNorm files provided by SomaLogic Inc. were bridged to data from the 1^st^ batch of samples using calibrators.

### Normalization of INTERVAL and LonGenity datasets

Relative Fluorescent Units (RFU) of each plasma protein were log10-transformed. We normalized the levels of each protein within each subcohort based on the average of the subjects in the 60-70 years range. Supplementary Fig. 11 shows representative normalization examples. Note that this normalization is needed when fitting aging trajectories (Fig. 2) but does not affect the results when “subcohort” is included as covariate in the modeling. The data from the 4 small independents cohorts were log10 transformed and bridged together using SomaLogic procedure based on calibrators. However, the number of samples in the 60-70 years range was too small to reliably bridge this data to the INTERVAL and LonGenity cohorts.

### Linear changes in the aging plasma proteome

To determine the effect of age and sex at the protein level, we used the following linear model:

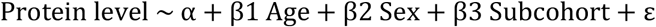

Type II sum of squares (SS) were calculated using the Anova function of the R car package^43^. This SS type tests for each main effect after the other main effects. Q-values were estimated using Benjamini–Hochberg approach^44^. It should be noted that the age range differs between cohorts. If the adjustment for cohort effect decreases the number of false positives, it could also alter the true positive rate. In the 4 small independent cohorts, the “Subcohort” covariate also accounts for batch effect since samples from different cohorts were measured in different batches (except for PRIN06 and GEHA that were measured together).

To determine the relative proportion of variance explained by age and sex, we calculated the partial Eta2 as follows:

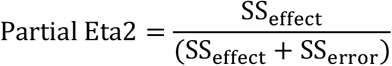

### Validation of the aging proteins

To provide confidence in the reproducibility of the protein assays, we compared our findings with the associations with age reported by Tanaka et al.^20^. To this end, we merged our results with those from Tanaka et al. using “SomaId”. Of note, Tanaka et al. used the same version of the SomaScan platform that we used for the 4 small independent cohorts (1,305 proteins).

### Sliding Enrichment Pathway Analysis (SEPA)

To determine the biological meaning of group of plasma proteins, we ranked the top 100 proteins based on the product of −log10(p-values) and beta age and queried three of the most comprehensive biological annotation and pathway databases: Gene ontology - GO^45^, Kyoto Encyclopedia of Genes and Genomes – KEGG^46^ and Reactome^47^. Using these databases, we tested enrichment for pathways in the top 10 to top 100 proteins in increments of 1 protein. The 2,925 proteins measured in this study cover 90% of the human GOs, reactome and KEGG terms containing more than 8 genes (Supplementary Fig. 12).

To analyze each incremental list of proteins, we used the R TopGO package^48^ for GO analysis and the R clusterprofiler package^49^ for KEGG and Reactome analyses. As input of SEPA, we used Gene Symbols provided by SomaLogic Inc. (Supplementary Table 1). The 2,925 proteins measured by SomaScan served as the background set of proteins against which to test for overrepresentation. Since several individual proteins (33 out of 2,925) were mapped to multiple Gene Symbols, we kept only the 1^st^ Gene Symbol provided by Somalogic to prevent false positive enrichment. For KEGG and Reactome analysis, clusterprofiler requires EntrezID as input. Therefore, we mapped Gene Symbols to EntrezID using the org.Hs.eg.db package^50^. Again, to avoid false positive enrichment, only the 1^st^ EntrezID was used when Gene Symbols were mapped to multiple EntrezID. Q-values were estimated using Benjamini–Hochberg approach^44^ for the different databases taken separately. For GO analysis, q-values were calculated for the three GOs classes (molecular function, cellular component, biological process) independently. To identify the most biologically meaningful terms and pathways, we reported only those with 20-500 proteins measured by the SomaScan assay. In addition, we focused on pathways consistently highly significant (q<0.05 for at least 20 different incremental list of proteins) and kept the top ten pathways per condition (e.g. for each wave of aging proteins). Ranking was performed based on the minimum fdr across the incremental lists of proteins. SEPA can be viewed as an extension of the GSEA approach^51^, with more control for true and false positives.

### Validation of the aging signature in mice

Male and virgin female C57BL/6JN mice were shipped from the National Institute on Aging colony at Charles River (housed at 67–73 °F) to the Veterinary Medical Unit (VMU; housed at 68–76 °F)) at the VA Palo Alto (VA). At both locations, mice were housed on a 12-h light/dark cycle and provided food and water ad libitum. The diet at Charles River was NIH-31, and Teklad 2918 at the VA VMU. Littermates were not recorded or tracked. Mice 18-months-old and younger were housed at the VA VMU for no longer than 2 weeks before euthanasia, and mice older than 18-months were housed at the VA VMU until they reached the experimental age. After anaesthetization with 2.5% v/v Avertin, blood was drawn via cardiac puncture. All animal care and procedures were carried out in accordance with institutional guidelines approved by the VA Palo Alto Committee on Animal Research.

Heterochronic parabiosis was conducted as previously described^6,13,52^ with 3- and 18-month-old mice. Briefly, incisions in the flank were made through the skin and peritoneal cavity of both mice, and adjacent peritoneal cavities were sutured together. Adjacent knee and elbow joints were then sutured together to facilitate coordinated locomotion. Skin was then stapled together using surgical autoclips (9-mm, Clay Adams), and mice were placed under heat lamps to recover from anesthesia. Each individual mouse was injected subcutaneously with Baytril antibiotic (5μg perg) and buprenorphine (0.05–0.1 μgml−1) for pain management, and 0.9% (w/v) NaCl for hydration. Mice were monitored and administrated drugs and saline over the next week as previously described.

EDTA-plasma was isolated by centrifugation at 1,000g for 10 min at 4 °C before aliquoting and storing at −80 °C. A total of 110 plasma samples were analyzed and aliquots of 150μl of plasma were sent on dry ice to SomaLogic Inc. (Boulder, Colorado, US). Samples were sent in two different batches, 29 samples in 2016 and 81 in 2018. Data for 1,305 SOMAmer probes were obtained and no sample or probe data were excluded. RFUs of each plasma protein were log10-transformed.

The SomaScan assay has been developed and validated for human fluids but successfully used in mouse research^9,23^. To understand how similar mouse and human sequences are, we downloaded all homologies between mouse and human along with sequence identifiers for each species (HOM_MouseHumanSequence.rpt) from MGI (http://www.informatics.jax.org/) as plain text files. Then, the protein reference sequences for both organisms were extracted from UniProt (https://www.uniprot.org/). On these matched sequence pairs, for each protein we computed a global pairwise sequence alignment. The alignments have been calculated by using the R “Biostrings” library^53^. The average identity was 0.85, supporting the use of the SomaScan assay with mouse plasma.

To determine the effect of age and sex at the protein level, we used the 81 samples from 1 month to 30 months. To this end, we fitted the following linear model:

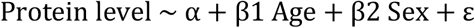

Type II sum of squares (SS) were calculated and Q-values were estimated using Benjamini–Hochberg approach.

To characterize the effects of young and old blood on the aging plasma proteome, normed scaled Principal Component Analysis (PCA) was performed using the R ade4^54^ package

### Prediction of human chronological age using the plasma proteome

To determine whether the plasma proteome could predict chronological age, we used glmnet^55^ and fitted a LASSO model (alpha=1, 100 lambda tested, “lamda.min” as the shrinkage variable estimated after 10-fold CV). Input variables consisted in Z-scaled log10 RFUs and sex information. Two-thirds (n=2,858) of the INTERVAL and LonGenity samples were used for training the model and the remaining 1,473 samples were used as a validation. In addition, the 171 samples from the 4 small independent cohorts were used to further assess the robustness of the predictive model.

To estimate whether a subset of the aging clock can provide similar predictive results, we used a two-step approach as we described previously^56^. One hundred models (100 lambda) including 0 to 373 proteins were created in step 1 and we estimated the accuracy of each of these model on the discovery and validation datasets, separately. Broken stick regression was used to determine the best compromise between number of variables and prediction accuracy.

### Associations between Δage and clinical and functions in old

We used the subjects from the LonGenity cohort to identify associations between deviations from the proteomic clock (Δage=predicted age-chronological age) and 334 clinical and functional variables (Supplementary Table 8). To this end, we tested the following linear model:

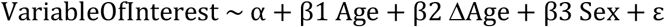

For binary outcomes, logistic regression was used. This analysis was separately performed in the discovery (n=2,858) and validation cohorts (n=1,473). Type II sum of squares were calculated using the Anova function of the R car package^43^.

### Clustering of protein trajectories

To estimate protein trajectories during aging, plasma proteins levels were z-scored and LOESS (locally estimated scatterplot smoothing) regression was fitted for each plasma factor. To group proteins with similar trajectories, pairwise differences between LOESS estimates were calculated based on the Euclidian distance and hierarchical clustering was performed using the complete method. To understand the biological functions of each cluster, we queried Reactome, KEGG and GO databases, as described above.

### Differential Expression - Sliding Window Analysis (DE-SWAN)

To identify and quantify linear and non-linear changes of the plasma proteome during aging, we developed a Differential Expression - Sliding Window ANalysis (DE-SWAN) approach. Considering, a vector l of k unique ages, we iteratively used l_k_ as the center of a 20-year window and compared protein levels of subjects in parcels above and below l_k_ i.e.[*l_k_* – 10y; *l_k_*[vs]*l_k_*; *l_k_* + 10y]. To test for differential expression, we used the following linear model:

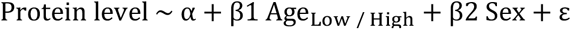

Age being binarized according to the parcels. For each l_k_, q-values were estimated using Benjamini–Hochberg correction. Type II sum of squares were calculated using the Anova function of the R car package^43^.

To assess the robustness and relevance of DE-SWAN results, we tested multiple parcel widths (5, 10, 15 and 20 years). In addition, we used multiple q-values thresholds and compared these results with those obtained by chance. To this end, we randomly permutated the phenotypes of the subjects and applied DE-SWAN to this new dataset. To keep the data structure, age and sex were permuted together. In addition, we analyzed the INTERVAL and LonGenity cohorts separately (Supplementary Fig. 6). Finally, we tested the same linear model when adjusting for Subcohort. This led to a loss of statistical power when the age range of the INTERVAL and LonGenity cohorts were overlapping but the three waves of aging proteins remained and the ranks of the top proteins were nearly identical (Supplementary Fig. 8). We used the model adjusted for subcohort when trying to understand the waves of aging proteins. The significance levels of the intersections between aging plasma protein signatures identified by linear modeling and DESWAN at different ages were determined using the R SuperExactTest package^57^.

### Relationships between the aging waves and the genome and the proteome of disease and traits

To quantify the overlap between proteins changing with age at different stages of life and the genome and the proteome of diseases and traits, we ranked DE-SWAN results based on p-values and created a k-ranked list of aging proteins, L_k_. To reflect the degree to which the genome/proteome are linked with the waves of aging plasma proteins, we walked down L_k_ and counted the number of proteins associated with the genome or specific proteome. When different versions of the SomaScan platform were used, we walked down L_k_ until reaching the top 100 proteins measured in both studies.

To identify specific genetic variants associated with the aging plasma proteome, we mined the summary statistics generated by Sun et al.^25^, who found 1,927 associations with 1,104 plasma proteins. Qgraph^58^ R package was used to create a network between the genome and the 2,925 proteins analyzed in this study.

To determine whether the aging proteome is associated with the proteome of clinical and functional variables, we used the subjects from the LonGenity cohort and tested the following linear model for the top variables identified in Supplementary Table 8:

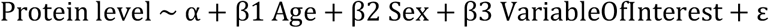

Type II sum of squares were calculated using the Anova function of the R car package^43^.

To determine whether the aging proteome is associated with disease proteomes, we integrated data and results from previous proteomic studies using the SomaScan platform. We re-analyzed one Alzheimer’s Disease - AD dataset publicly available by AddNeuroMed^26^ and used summary statistics from published studies focused cardiovascular disease - CVD^28^, Down Syndrom - DS^27^ and body mass index - BMI^29^.

AddNeuroMed is a European multi-center study in which the AD proteome was quantified in plasma samples from 681 controls, Mild Cognitive Impairment (MCI), and AD subjects using a previous version of the SomaScan assay. Files used were downloaded from the Synapse portal in March 2016 (syn5367752) and included measurements of 1016 plasma proteins from 931 samples. We limited our analysis to the 645 samples available at visit 1 (191 control, 165 MCI and 289 AD). Raw data were log10-transformed. Four samples (2 control and 2 AD) were considered as outliers based on principal component analysis and filtered out.

To identify plasma proteins associated with AD, we used linear models with diagnosis, age, sex and center as covariates:

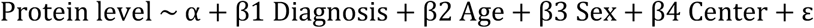

Type II sum of squares were calculated using the Anova function of the R car package^43^. To determine whether aging plasma proteins are involved in other disease signatures, we identified three studies using the SomaScan platform in large human cohorts providing detailed summary statistics. Carayol et al. mined the plasma proteome to obtain new insights into the molecular mechanisms of obesity. Out of 1,129 proteins measured, they identified 192 plasma proteins significantly associated with BMI (p<0.05 after Bonferroni correction). Summary statistics we used were obtained from their “Supplementary Data 1”^29^. Ganz et al. derived and validated a 9 protein risk-score to predict risk of cardiovascular outcomes^28^. In addition to these 9 proteins, 191 other proteins were significantly associated with cardiovascular risk (p<0.05 after Bonferroni correction). Summary statistics for these 200 proteins are available in their eTable 4 and the 1,130 proteins measured of this study are listed in their eTable 1. Finally, Sullivan et al. used an extended version of the SomaScan platform to study DS and identified a large number of dysregulated proteins^27^. We used the results for the Discovery cohort (sheet A of the “Supplementary File 1”) in which, 258 proteins out of 3,586 proteins are reported to be associated with DS (p<0.05 after Bonferroni correction).

Since detailed protein information was not available for all studies, we used either gene symbols or unitprotID to merge disease proteomes characterized in published studies with the aging proteome identified in this study. When multiple p-values were reported for the same gene symbol (or a combination of gene symbols), only the most significant p-value was retained.

### Code availability

An R package for DE-SWAN is available in github: https://github.com/lehallib/DEswan

### Data availability

We created a searchable web interface to mine the human INTERVAL and LonGenity datasets (https://twc-stanford.shinyapps.io/aging_plasma_proteome/)

The Human independent cohorts and mouse protein data are available in Supplementary Tables 16 and 17. The Interval data is available through the European Genome-Phenome Archive (https://ega-archive.org/studies/EGAS00001002555).

---

**Supplementary Figure 1:**
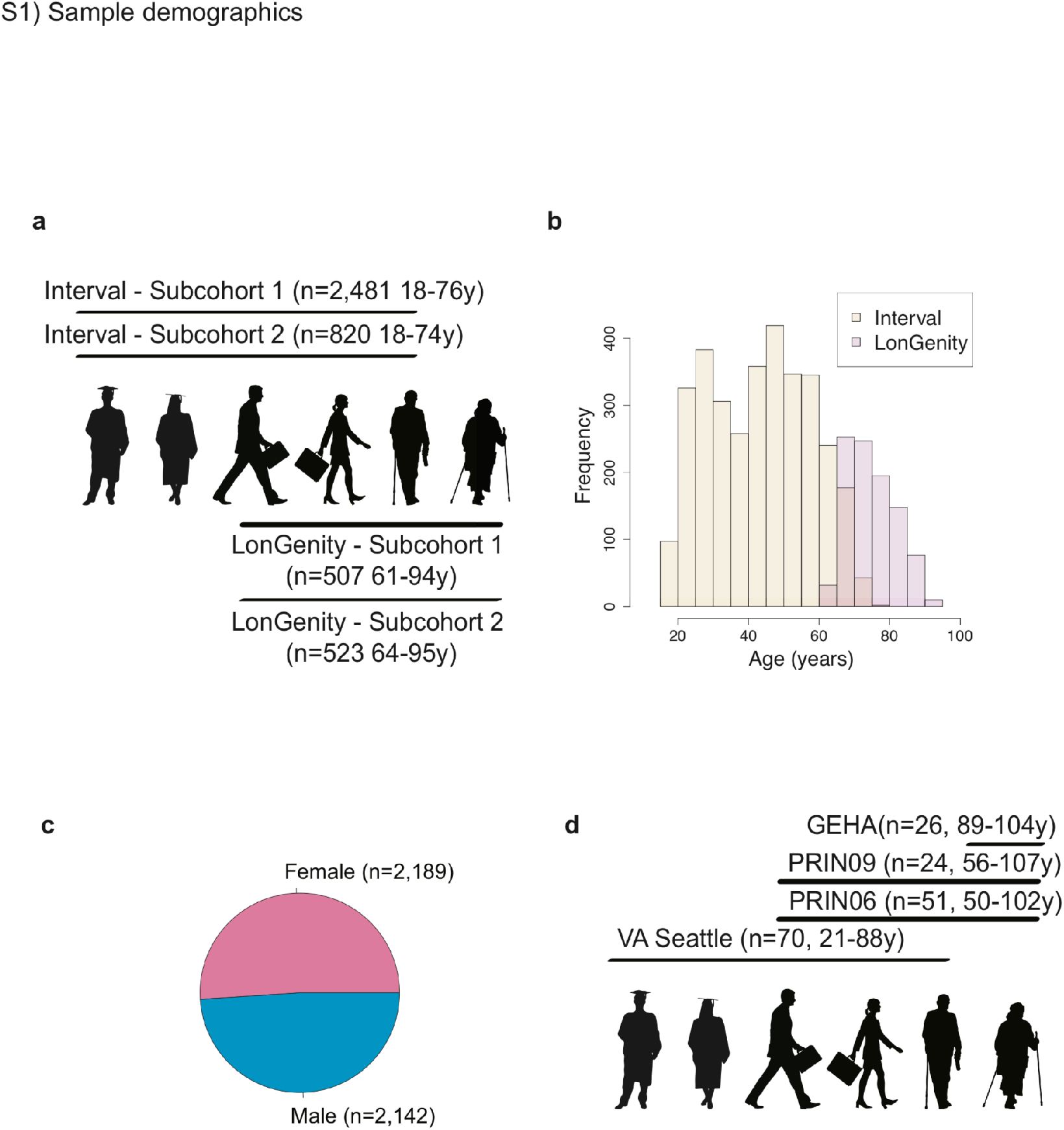
Sample demographics. Age (a, b), cohort (a, b) and sex distributions (c) of the 4,331 subjects from the Interval and LonGenity cohorts. (d) Age and cohort distributions of the 171 subjects from the 4 independent cohorts.

**Supplementary Figure 2:**
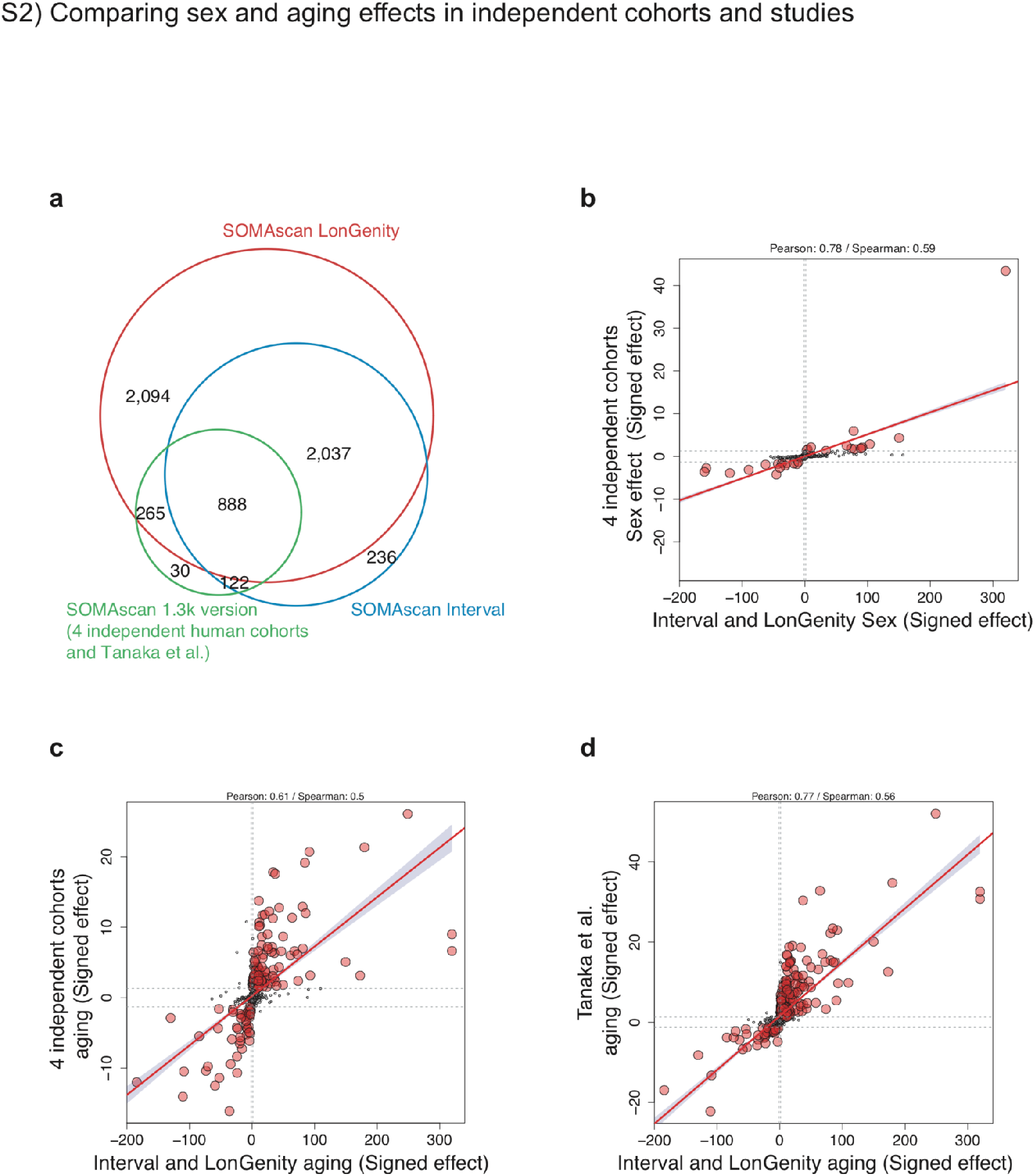
Comparing aging effect in independent cohorts and studies. (a) Age and sex effects in the INTERVAL and LonGenity studies were compared to age and sex effects in 4 independent cohorts analyzed together and to age effect from Tanaka et al. (2018) The aging plasma proteome was measured with the SomaScan assay in these cohorts and 888 proteins were measured in all studies (b) Scatter plot representing the signed −log10(q value) of the sex effect in the INTERVAL/LonGenity cohorts (x axis) vs the 4 independent cohorts (y-axis). Similar analysis for the age effect in the 4 independent cohorts (c) and in Tanaka et al (2018) study (d)

**Supplementary Figure 3:**
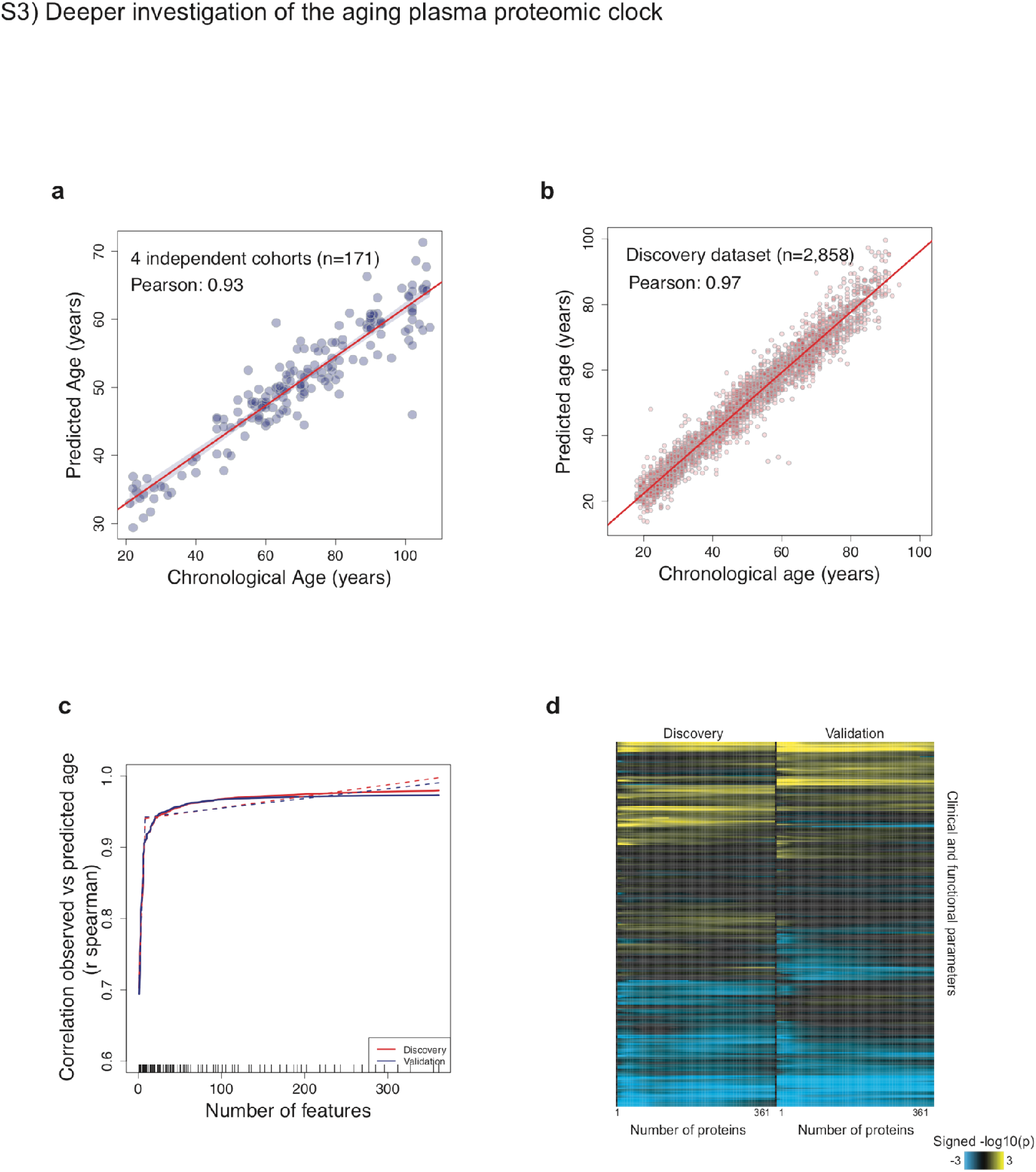
Deeper investigation of the aging proteomic clock. (a) Prediction of age in the 4 independent cohorts (n=171) using the proteomic clock. Only 141 proteins out of the 373 constituting the clock were measured in these samples. (b) Prediction of age in the discovery cohort (n=2,858) using 373 plasma markers. (c) Feature reduction of the aging model. A two-step approach was used to estimate whether a subset of the aging signature can provide similar results to the 373 aging proteins. The proteins were ranked according to the absolute value of their coefficients in the LASSO model. Then a ridge regression model was built using the discovery dataset with the top 2 variables selected in step 1. This process was then repeated iteratively using the top 3, 4, […] up to top 373 variables selected in the first step. Dashed lines represent a broken stick model and indicate the best compromise between number of variables and prediction accuracy. (d) Heatmap representing the associations between delta age and 334 clinical and functional variables. As in (c) the analysis was performed for the top 2 to top 373 variables predicting chronological age. The non-uniformity in the heatmaps suggests that specific subsets of proteins may best predict certain clinical and functional parameters.

**Supplementary Figure 4:**
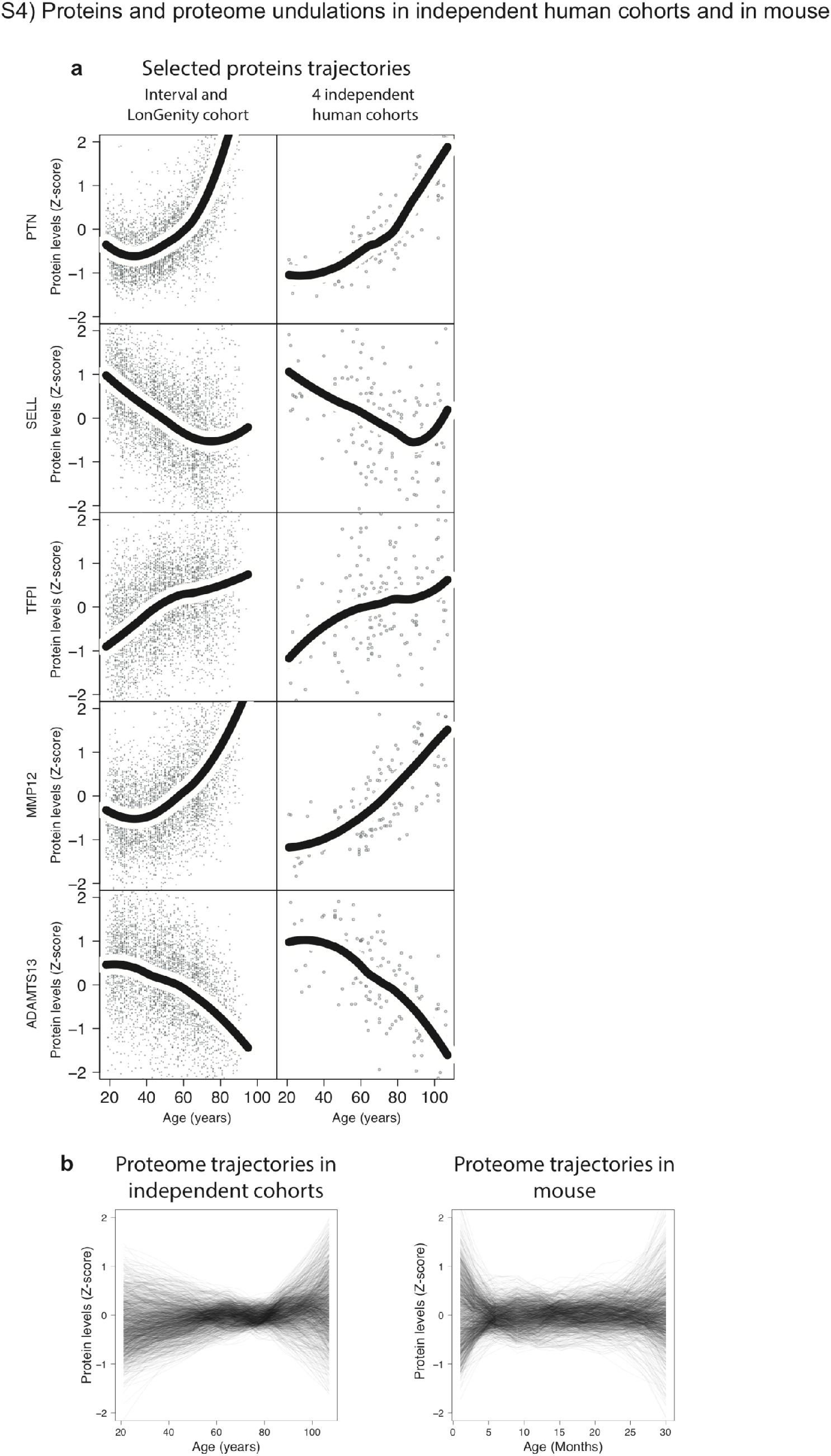
Proteins and proteome undulations in independent human cohorts and in mouse. (a) Trajectories of 5 selected proteins based on the Interval and LonGenity cohorts (n=4,331, left) and 4 independent human cohorts (n=171, right). Trajectories were estimated using LOESS regression. (b) Undulation of the 1,305 plasma proteins in 4 independent cohorts (n=171, left) and in mouse (n=81, right). Plasma proteins levels were z-scored and LOESS regression was fitted for each plasma factor.

**Supplementary Figure 5:**
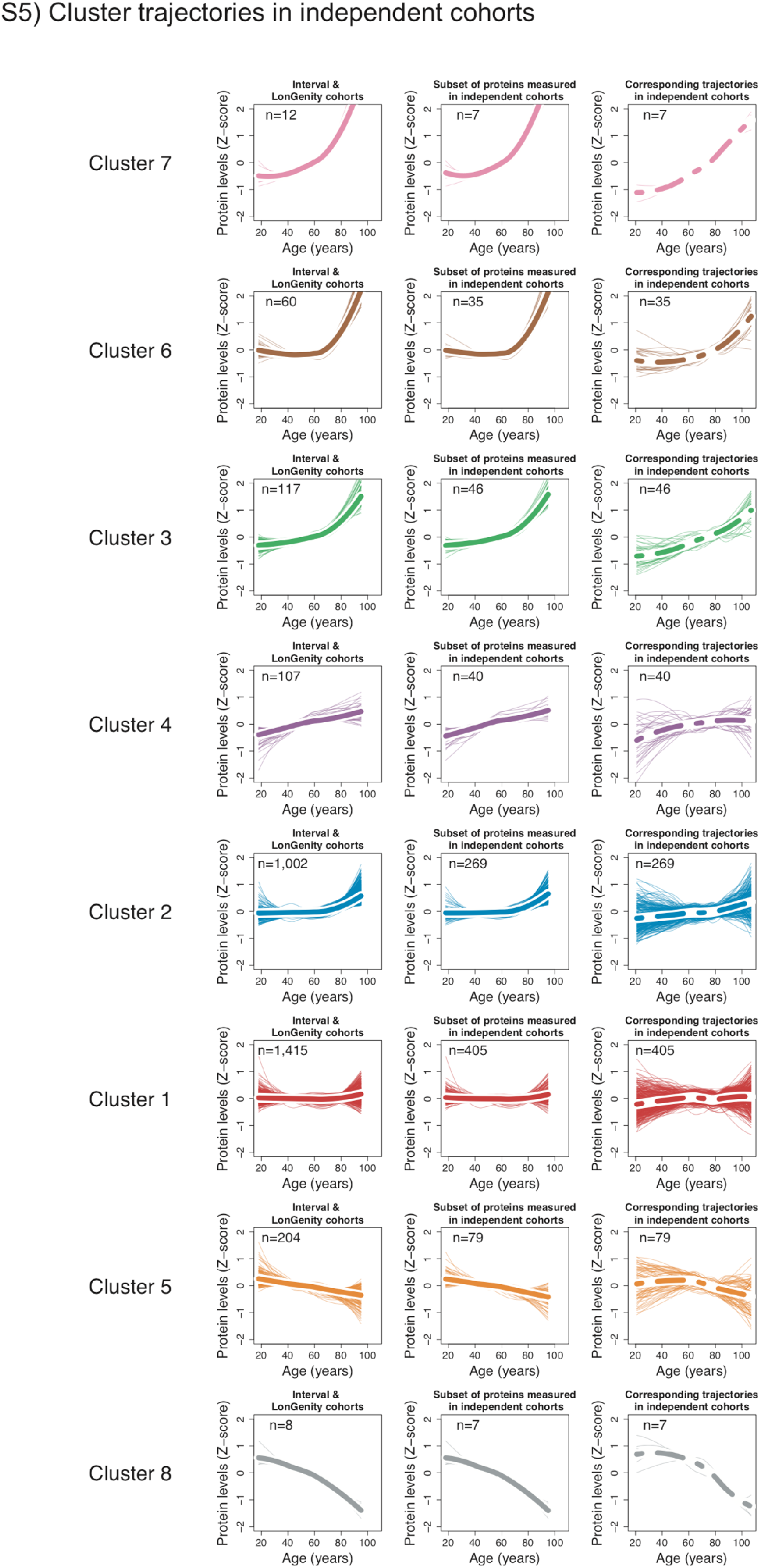
Cluster trajectories in independent cohorts. Protein trajectories for the 8 clusters identified in the Interval and LonGenity cohorts (left column). Thicker lines represent the average trajectory for each cluster. Cluster trajectories for the subset of proteins measured in the 4 independent cohorts (middle column). Corresponding cluster trajectories in 4 independent cohorts (right column).S6) Top pathways in clusters

**Supplementary Figure 6:**
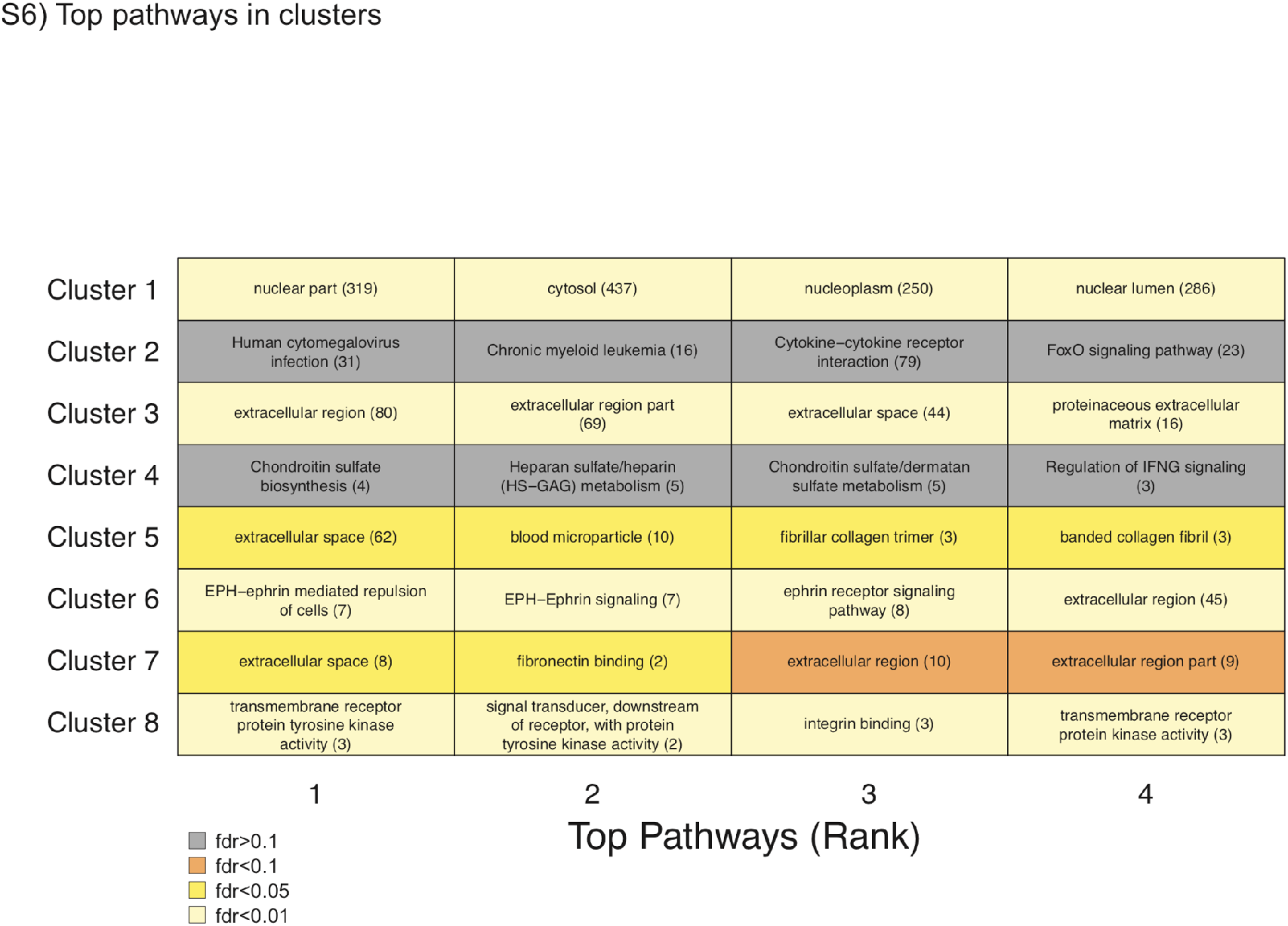
Pathway enrichment was tested using GO, Reactome and KEGG databases. The top 4 pathways for each cluster are shown. Pathway IDs and number of plasma proteins associated are represented in the table.

**Supplementary Figure 7:**
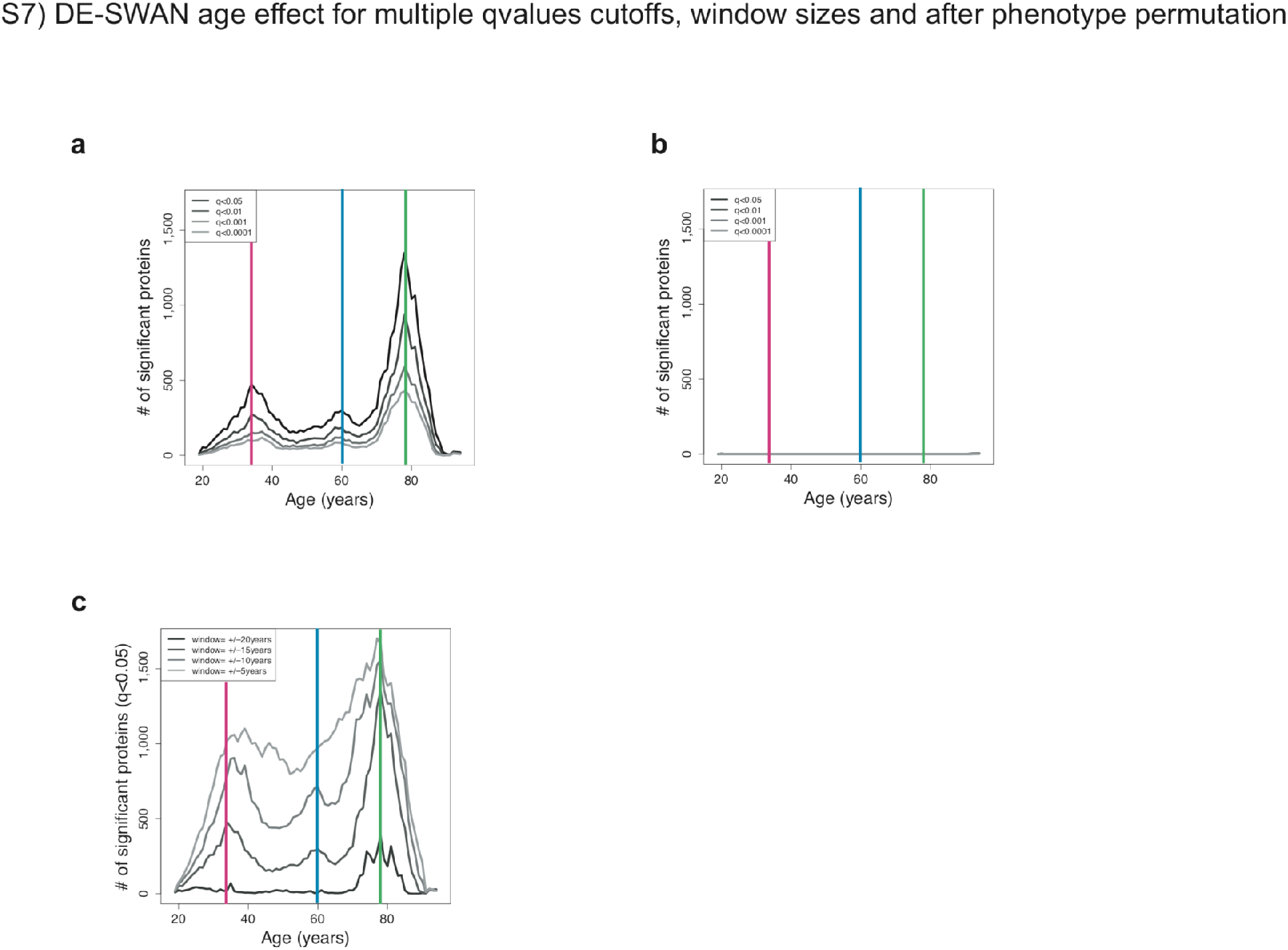
DE-SWAN age effect for multiple q-values cutoffs, windows size and after phenotypes permutations. Different Q-value cutoffs are represented in (a). Similar analysis with different after phenotype permutations (b). and different windows size in (c). The 3 local peaks identified at age 34, 60 and 78 are indicated by colored vertical lines.

**Supplementary Figure 8:**
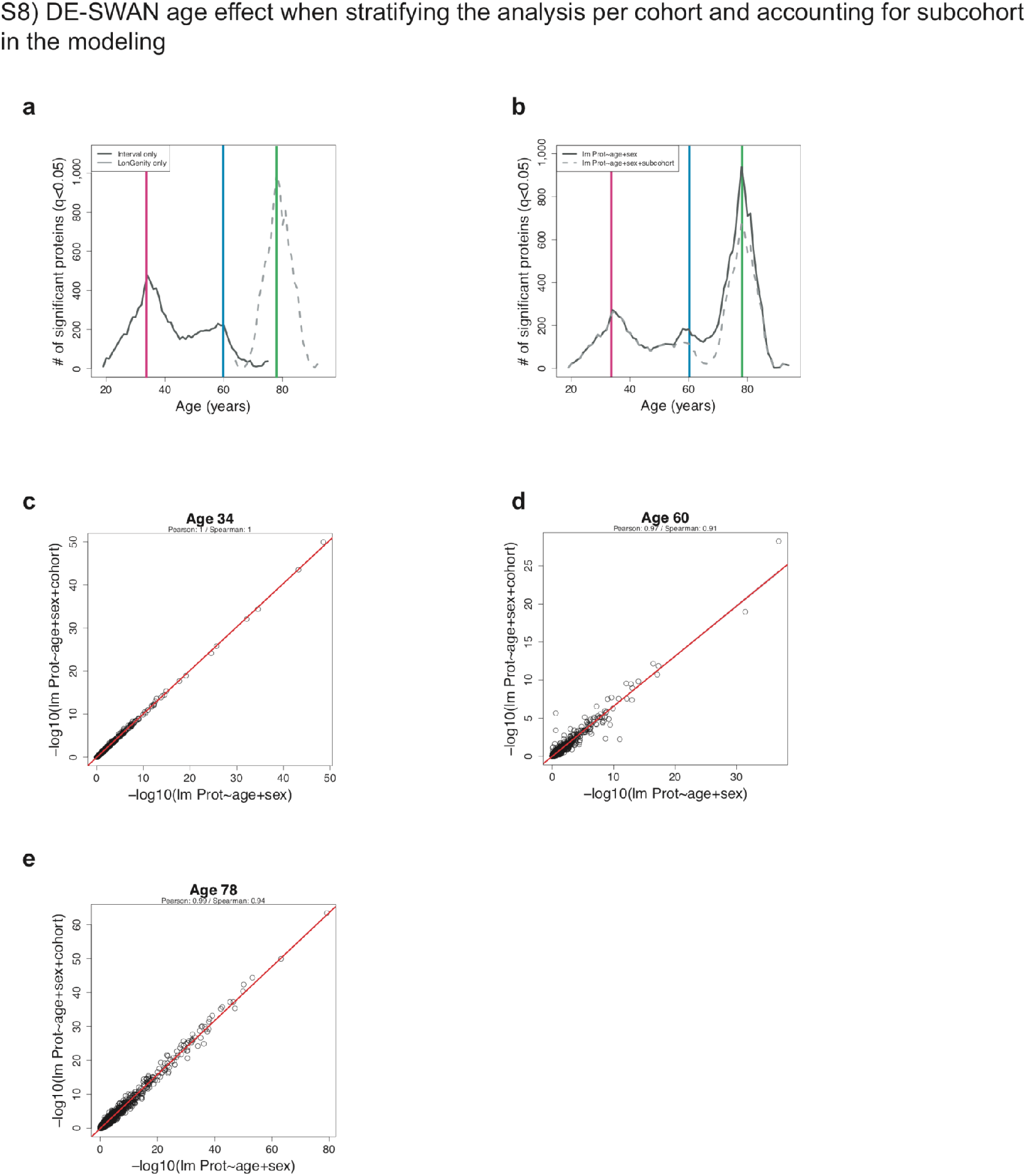
DE-SWAN age effect when stratifying the analysis per cohort and accounting for subcohort. (a) DE-SWAN results when stratifying the analysis per cohort. The 3 local peaks identified at age 34, 60 and 78y are indicated by colored vertical lines. (b) Comparison of DE-SWAN results when adjusting for cohort or not. (c-e) Age effect at 34, 60 and 78y when adjusting for cohort or not.

**Supplementary Figure 9:**
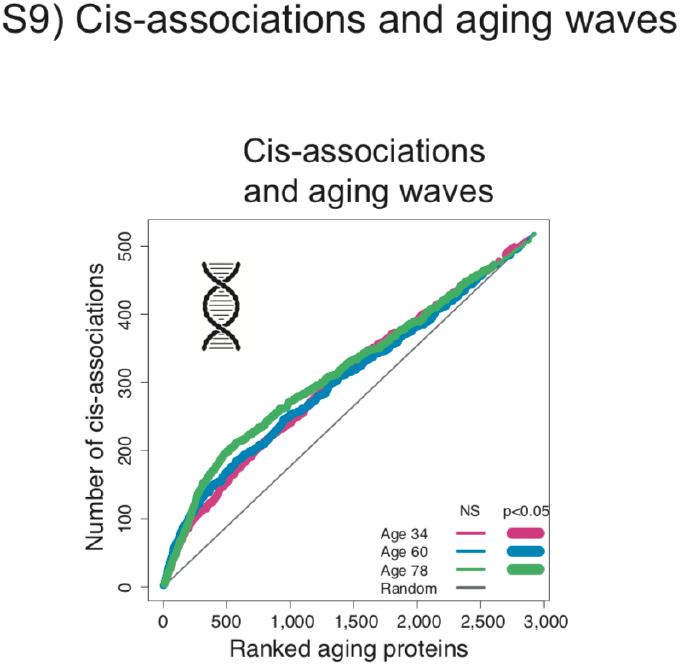
Cis-associations and aging waves. Enrichment for cis-association in the waves of aging proteins identified by DE-SWAN. Aging proteins were ranked based on p-values at age 34, 60 and 78 and the cumulative number of cis-associations was counted. Permutation based tests (1e+5) were used to assess significance.

**Supplementary Figure 10:**
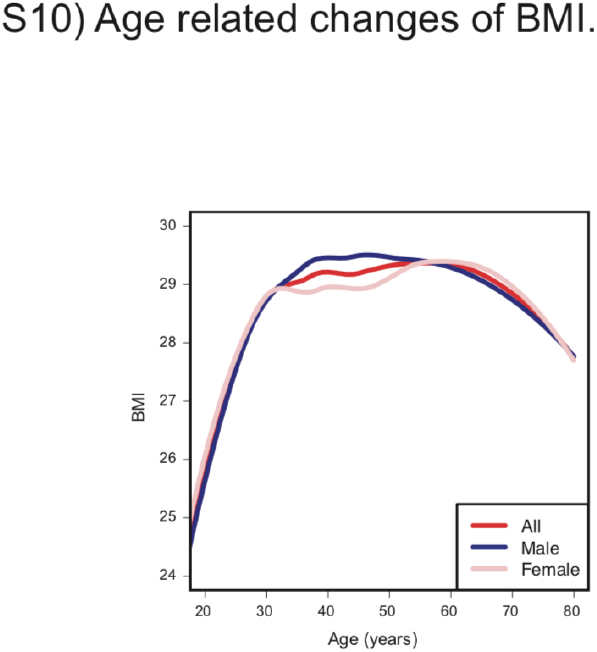
Age related changes of BMI. Changes during aging are estimated using LOESS curves, fitted for all subjects and for male and female separately. Data from the R NANHES package.

**Supplementary Figure 11:**
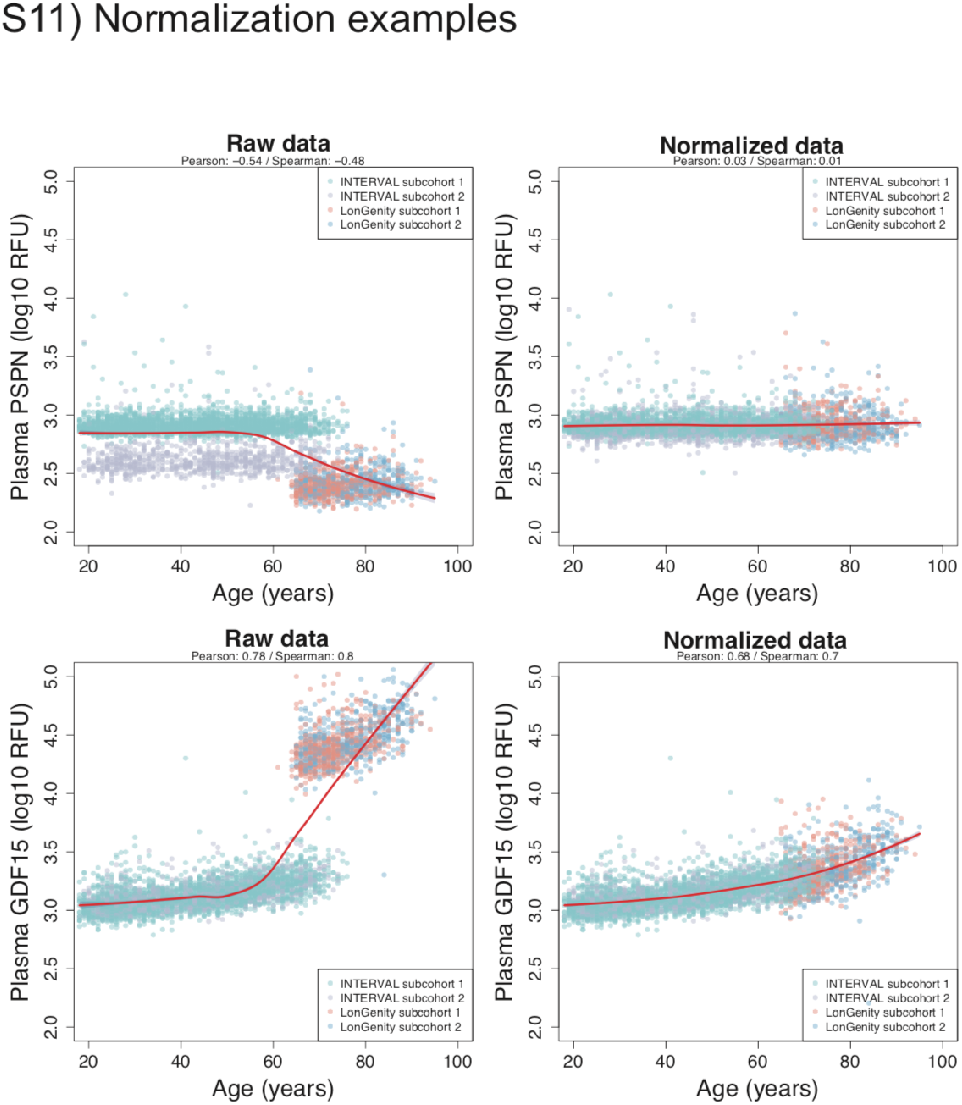
Normalization examples. Representative examples of the normalization process in the Interval and LonGenity cohorts. For each protein, the average of the subjects in the 60-70y range within each subcohort was used as a normalization factor. LOESS regression curves are fitted. Note that this normalization is needed when fitting aging trajectories (Fig. 2) but does not affect the results when “subcohort” is included as covariate in the statistical modeling.

**Supplementary Figure 12:**
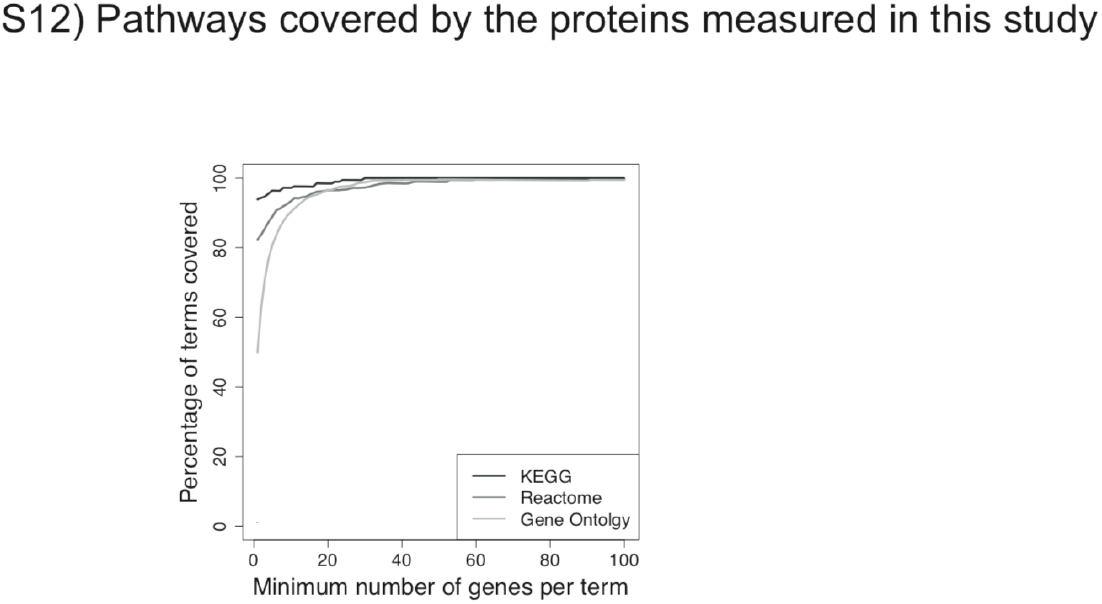
Pathways covered by the 2,925 proteins measured in this study. The 2,925 proteins measured in this study cover 90% of the human GO, Reactome and KEGG terms containing more than 8 genes. Coverage for pathways with 1+ to 100+ genes is represented.

